# Hidden limbs in the “limbless skink” *Brachymeles lukbani:* developmental observations

**DOI:** 10.1101/2020.12.09.418483

**Authors:** Daniel Smith-Paredes, Oliver Griffith, Matteo Fabbri, Laurel Yohe, Daniel G. Blackburn, Cameron D. Siler, Bhart-Anjan S. Bhullar, Günter P. Wagner

## Abstract

Reduced limbs and limblessness have evolved independently in many lizard clades. Skinks exhibit a wide range of limb-reduced morphologies, but only some species have been used to study the embryology of limb reduction (i.g., digit reduction in *Chalcides* and limb reduction in *Scelotes*). The genus *Brachymeles*, a Southeast Asian clade of skinks, includes species with a range of limb morphologies, from pentadactyl to functionally as well as structurally limbless species. Adults of the small, snake-like species *Brachymeles lukbani* show no sign of external limbs in the adult except for small depressions where they might be expected to occur. Embryos of *B. lukbani* in early stages of development, on the other hand, show a truncated but well-developed limb with a stylopod and a zeugopod, but no signs of an autopod. As development proceeds, the limb’s small size persists even while the embryo elongates. These observations are made based on external morphology. We used florescent whole-mount immunofluorescence to visualize the morphology of skeletal elements and muscles within the embryonic limb of *B. lukabni*. Early stages have a humerus and separated ulna and radius cartilages; associated with these structures are dorsal and ventral muscle masses as those found in the embryos of other limbed species. While the limb remains small, the pectoral girdle grows in proportion to the rest of the body, with well-developed skeletal elements and their associated muscles. In later stages of development, the small limb is still present under the skin but there are few indications of its presence, save for the morphology of the scale covering it. The adult morphology consists of a well-developed pectoral girdle, small humerus, extremely reduced ulna and radius, and well-developed limb musculature connected to the pectoral girdle. These muscles form in association with a developing limb during embryonic stages, a hint that “limbless” lizards that possesses these muscles may have or have had at least transient developing limbs, as we find in *B. lukbani.* Overall, the observed pattern of ontogenetic reduction, leading to an externally limbless adult in which a limb rudiment is hidden and covered under the trunk skin, is a situation called cryptomelia. The results of this work add to our growing understanding of clade-specific patterns of limb reduction and the convergent evolution of limbless phenotypes through different developmental processes.

## 1 INTRODUCTION

Limb reduction and limblessness have evolved many times independently within squamate reptiles (Gans, 1975; Greer, 1991). Snakes are easily the most recognizable limbless clade. Although some groups of snakes (Aniliidae, Boidae, Leptotyphlopidae, Pythonidae, Typhlopidae; List, 1966) possess rudimental hindlimbs, with pelvic and proximal limb skeletal elements, forelimbs and pectoral girdles are not present in any extant species. Similarly, almost every large clade of lizards has, in fact, evolved its own snake-like morphotype at least once, including Amphisbaenidae, Anguidae, Cordylidae, Dibamidae, Gekkota, Gymnophthalmidae, Gerrhosauridae, and Scincidae (Greer, 1991; Leal and Cohn, 2018). Limbs could, in theory, be reduced or absent as a consequence of a variety of developmental mechanisms such as extreme allometry, degeneration and agenesis. Among the various cases of convergence towards limbless body plans among clades of lizards, embryonic development has been investigated in only a handful of lineages, all of which are species of snakes or anguid and scincid lizards (Raynaud, 1985; Infante *et al.*, 2018), and as a consequence our understanding of similarities and differences in the ontogeny of limb loss is limited. In python embryos, hindlimbs develop but have no apical ectodermal ridge (AER), nor expression of genes normally associated with maintenance of limb growth (Cohn and Tickle, 1999). Hindlimb development becomes truncated and at least some distal skeletal elements fail to form. In contrast, forelimb development is never initiated (Cohn and Tickle, 1999). In snakes lacking hindlimb remnants that have been studied (Zehr, 1962; Raynaud, 1985; Jackson, 2002), neither fore- nor hindlimbs initiate development. The developmental pattern of the reduced hindlimb of python embryos differs from that of other limbless lizards (Raynaud, 1985). In comparison, in a study of the limbless anguid genus *Anguis,* fore- and hindlimb rudiments appear in early stages; however, development soon ceases and regression and sequential disappearance of the forelimb and then the hindlimb occurs (Raynaud, 1985). This pattern is similar to observations made in another extremely limb reduced anguid, *Pseudopus* (referred to as *Ophisaurus* in the references), in which fore- and hindlimb buds also start to develop before subsequent degeneration and disappearance occurs (Rahmanl, 1974; Raynaud, 1985).

Skinks of the genus *Scelotes* show different degrees of limb reduction, including limbless forms (Lande, 1978; Wiens and Slingluff, 2001; Siler and Brown, 2011). Embryos of different species form both fore- and hindlimb buds, which stop developing and regress to different degrees (Raynaud *et al*., 1975; Raynaud and Van den Elzen, 1976; Raynaud, 1985). For example, embryos of *Scelotes inornata* form a rudimentary AER which later degenerates differentially among the fore- and hindlimbs (Raynaud, 1985). In the forelimb, regression occurs rapidly, while in the hindlimb it does so more slowly, resulting in an adult with a rudimentary hindlimb possessing a proximal portion of the femur (Raynaud, 1985). In *S. brevipes*, limb development follows a similar pattern; however, the hindlimb develops further and the adult retains an ossified femur and fused cartilaginous tibia and fibula (Raynaud, 1985). Finally, *S. gronovii* embryos have a well-developed AER in the hindlimb, and the adult hindlimb has an ossified femur, tibia, fibula, and one finger with three phalanges, while the AER is not well developed in the forelimb, and degenerates early in development (Raynaud, 1985).

Among squamate clades, by far the greatest diversity of independent origins of limb reduction and limblessness occurs in skinks (family Scincidae), which have evolved limb reduced forms more times than any other lizard group (Greer, 1991; Russell and Bauer, 2008). Furthermore, a number of skink genera include suites of closely related species that display the full spectrum of body forms, from pentadactyl to limbless, including *Brachymeles* (Wagner *et al.*, 2018; Siler and Brown, 2011), *Chalcides* (Carranza *et al*., 2008; Young *et al.*, 2009), *Lerista* (Skinner *et al.*, 2008; Skinner and Lee, 2009) and *Scelotes* (Raynaud, 1985), making them attractive model clades for studying evolutionary convergence in phenotype and major transitions in body form. In this study, we investigate the anatomy of embryos of *Brachymeles lukbani*, a recently described, elongated, slender skink without any trace of external limbs in adults, except for a small depression where the limb could be expected to be found (Siler *et al.*, 2010). Our results provide new information on the developmental patterns leading to the origin of limblessness and clues into the sequence of evolutionary events behind the evolution of repeated limb reduction and loss in lizards.

## 2 METHODS

Embryos of *Brachymeles lukbani* were collected in the field during an expedition to the Philippines in May and June 2016. Surveys for individuals of *B. lukbani* were conducted at Mt. Labo, Barangay Tulay Na Lupa, Municipality of Labo, Camarines Norte Province, Luzon Island, in coordination with local community partners. Animals were captured by hand raking leaf litter and loose soil surrounding tree root networks and rotting logs along the forest floor (Siler and Brown, 2011). Pregnant female individuals were euthanized and prepared as vouchered specimens after embryos were extracted for subsequent preparation (Simmons, 2015). Vouchered specimens were deposited in the National Museum of the Philippines and the Sam Noble Oklahoma Museum of Natural History.

Corn snake (*Pantherophis guttatus*) eggs were obtained from a colony housed at Trinity College. Embryos were staged according to (Zehr, 1962), collected and dissected in cold PBS, then fixed in 4% PFA in a shaker at 4°C for 7 days, then dehydrated 3–4 times with 15 minutes washes of Methanol 100% and stored at −20°C until further processing.

Dehydrated embryos were bleached overnight in a solution of Methanol:DMSO:H_2_O_2_ 4:1:1 under light. After bleaching they were washed with Methanol 100% two times for ten minutes and then rehydrated in increasing concentrations of PBS:Methanol (25%,50%,75%,100%). After two extra washes in PBS, embryos were placed in a solution of 4% Acrylamide in PBS, with 0.25% VG44 as initiator al left at 4°C overnight. Next day, embryos were placed in a 50 mL falcon tube with a special adaptor, and O_2_ was replaced by N_2_ by taking out air with a vacuum chamber and pumping N_2_ from a tank. Embryos were incubated at 37°C for 4 hours to allow acrylamide to polymerize, and later were washed in a solution of 200mM SDS 200mM Boric Acid in distilled water until they became transparent. When transparent, embryos were washed for an hour six times in PBS with 1% TritonX-100 (PBSt). Immunostaining was performed using two antibodies targeting myosin heavy chain (MF-20, DSHB) and the transcription factor Sox9 (Sox9, AB5535, Abcam) in concentrations 1:50 and 1:1000 respectively, in a solution with 5% DMSO, 5% normal horse serum in PBSt. Antibodies were incubated overnight, washed six times for one hour in PBSt, and incubated with secondary antibodies (Goat anti-mouse 555, Goat anti-rabbit 647 Invitrogen) overnight. Embryos were then washed in PBSt three times and stored in RIMS (Refractive Index Matching Solution; Yang *et al.*, 2014).

Embryos stored in RIMS were either photographed directly in RIMS or accommodated in liquid agarose + RIMS (1% low temperature melt Agarose GPG/LMP, AmericanBio, dissolved in RIMS). Embryos were imaged with a Zeiss Axio Zoom.V16 fluorescent scope or with a Zeiss LSM880 Confocal Microscope collecting multiple tiles of Z-stacks, according to the size of the embryo. 3D projections of images were reconstructed using Fiji software (Schindelin *et al.*, 2012).

Two adult *B. lukbani* specimens were stained in 5% I_2_KI for 15 days, fixed in agarose 1%, and mounted for scanning in a 50 mL tube. The specimens were scanned on a high-resolution Nikon H225 ST μCT-scanner at Yale University. Scan parameters included 0.00967746 mm voxel size resolution, 105 kV, 64 μA, and centered at a region focused on the head and forelimb. The scan image stacks were imported into in-house Nikon post-scanning image processing software, where they were reconstructed with dual high-resolution centers of rotation and 3^rd^ level beam hardening. Resulting image stacks were imported into VGStudio Max v. 3.4.1 for segmentation.

## 3 RESULTS

### 3.1 External embryonic morphology of *Brachymeles lukbani*

The youngest embryo of *B. lukbani* (Figure 1, OMNH 45693) has a slightly elongated morphology, with a long body coiled once. The heart lies still outside of the thoracic cavity. The forelimb is small but almost complete, with a bent elbow between the stylopod and zeugopod. No autopod seems to be present, as there is no evident digital paddle. In the second embryo (Figure 1, OMNH 45709), differential growth has resulted in a more elongated body shape, and a smaller looking limb. The heart is now enclosed in the thoracic cavity. The limb looks proportionally smaller, in relation to the rest of the body and the bent elbow is less obvious. In later stage embryos (Figure 1, OMNH 45717, 45760), eyelids have started to cover the eye, scales have developed and are pigmented, and the limb is limited to a small protuberance on the side of the body, covered by a small, rounded scale.

**FIGURE 1.**
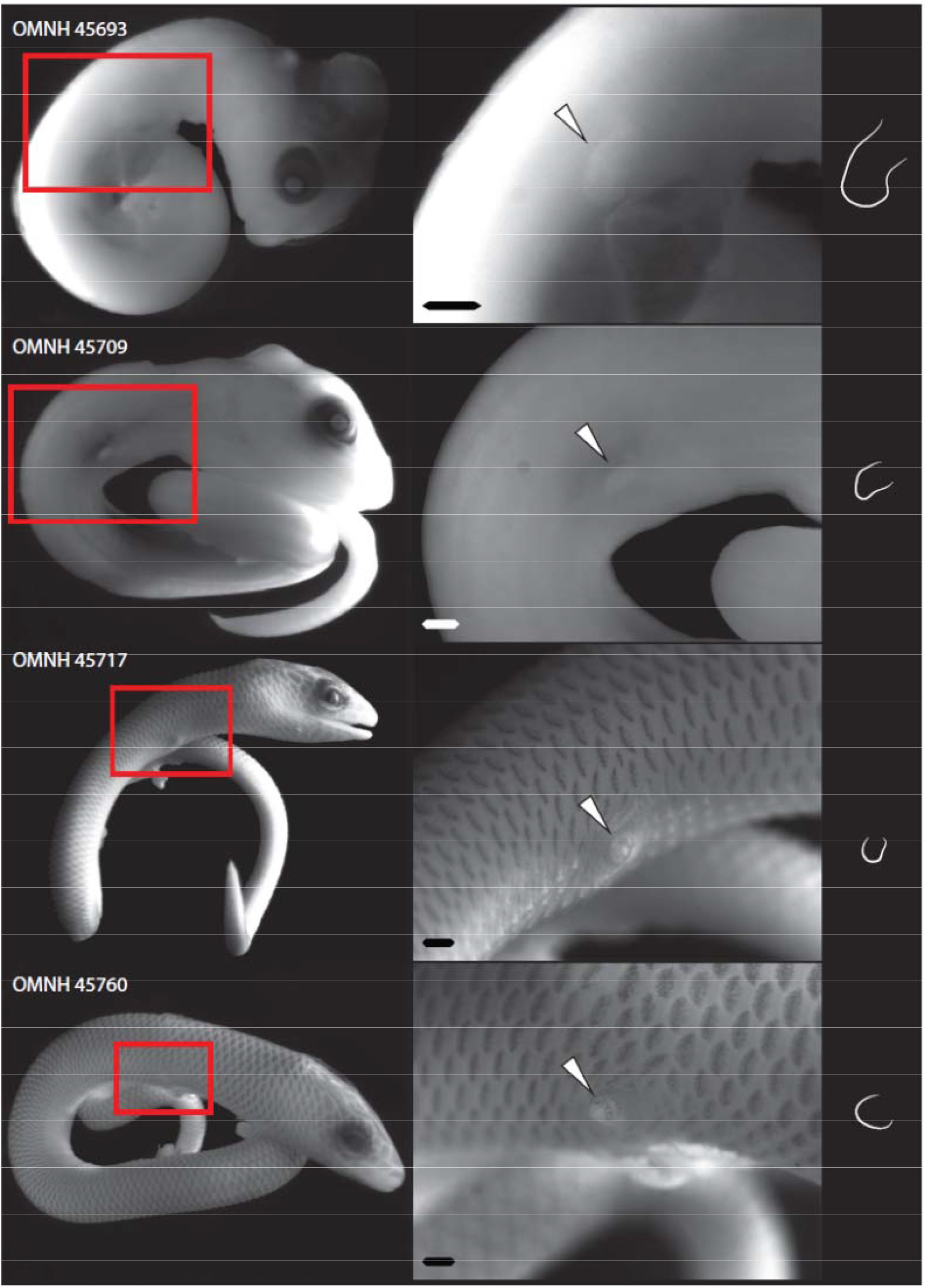
The external morphology of a series of embryos of *Brachymeles lukbani.* Left column shows the whole embryo, middle column shows a close up on the forelimb, right column shows a line drawing of the shape of the forelimb. From earlier to later stages, the limb does not grow considerably and gets covered with small scales. Scale bar: 500 μm

### 3.2 Skeletal embryonic development of *Brachymeles lukbani*

The youngest of the *Brachymeles lukbani* embryos is in an early stage of skeletal development. As revealed by immunostaining against Sox9 protein, to label pre-cartilaginous and early cartilaginous condensations, vertebrae and chondrocranial components have already started to develop (Figure 2, OMNH 45693). In the pectoral region, the developing scapulocoracoid plate can be seen as a continuous structure, and a small humerus, ulna, and radius are present in the arm.

**FIGURE 2.**
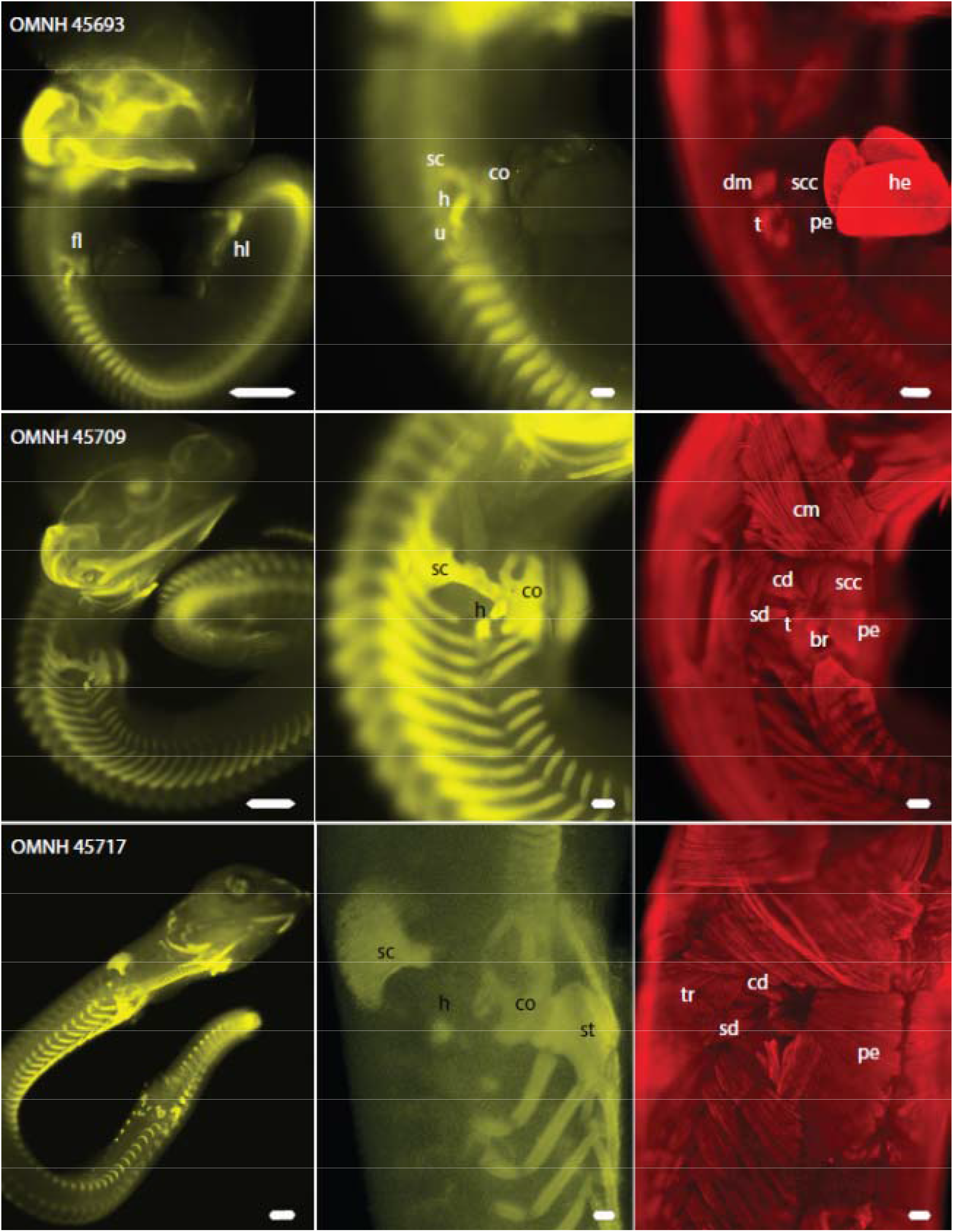
Musculoskeletal anatomy of the developing embryos of *Brachymeles lukbani* visualized with immunofluorescence. Left and middle columns show Sox9 labelled in yellow, right column shows Myosin heavy chain labelled in red. br: brachial musculature, co: coracoid plate, cd: clavicular deltoid muscle, cm: cleidomastoid muscle, dm: deltoid musculature, fl: forelimb, h: humerus, he: heart, hl: hindlimb, pe: pectoral muscle, sc: scapular plate, scc: supracoracoid muscle, sd: scapular deltoid muscle, st: sternum, t: triceps musculature, tr: trapezius muscle, u: ulna

In the next stage of development (Figure 2, OMNH 45709), rib precursors are visible as extended projections off of the vertebral condensations. In the skull, the chondrocranial elements are more differentiated, with distinct quadrate, optic and nasal capsules, and a well-defined hyoid apparatus. The scapular plate and the coracoid plate are differentiated, including the excavations that will form the margin of the fenestrae typical of lizard primary girdles. The humerus is much longer than the ulna and radius, and the label seems to be interrupted in the diaphysis, which could be a sign of cartilage maturation. The ulna and radius are in close contact and no other skeletal element has developed distally.

In later stages (Figure 2, OMNH 45717) maturation of cartilage has proceeded, as evidenced by the weak or absent Sox9 signal in portions of the ribs, vertebrae, and elements of the chondrocranium and Meckel’s cartilage. Ribs and tracheal rings are still Sox9 positive, as are portions of the suprascapular, coracoids, and presternum. There is no evidence of Sox9 positive cells in the forelimb skeleton.

### 3.3 Muscular embryonic development in the forelimb of *Brachymeles lukbani*

In amniotes, premuscular cells of somitic origin invade the limb and form dorsal and ventral muscle masses, flanking the skeletal condensations. These masses later divide into individual muscles of the chest, shoulder, arm, and hand (Romer, 1944; Christ and Brand-Saberi, 2004). In the earliest embryo observed (Figure 2, OMNH 45693), the division of these muscle masses has already started. The dorsal mass is split into identifiable Deltoid, Latissimus, Triceps, and forelimb Extensor divisions, while ventrally a Pectoral, Supracoracoideus, Biceps, and forelimb Flexor divisions are apparent. In the next stage (Figures 2 and 3, OMNH 45709), the shoulder and chest muscle masses, extrinsic to the arm, have divided into identifiable individual muscles, although they are small in comparison to the axial muscles in the region. The intrinsic muscles of the arm seem to be much less developed, possibly even degenerating. Later stages (Figure 2, OMNH 45717) do not show any evidence of intrinsic arm musculature, but small extrinsic arm muscles (Deltoid, Latissimus, Pectoral, Supracoracoid musculature) remain.

**FIGURE 3.**
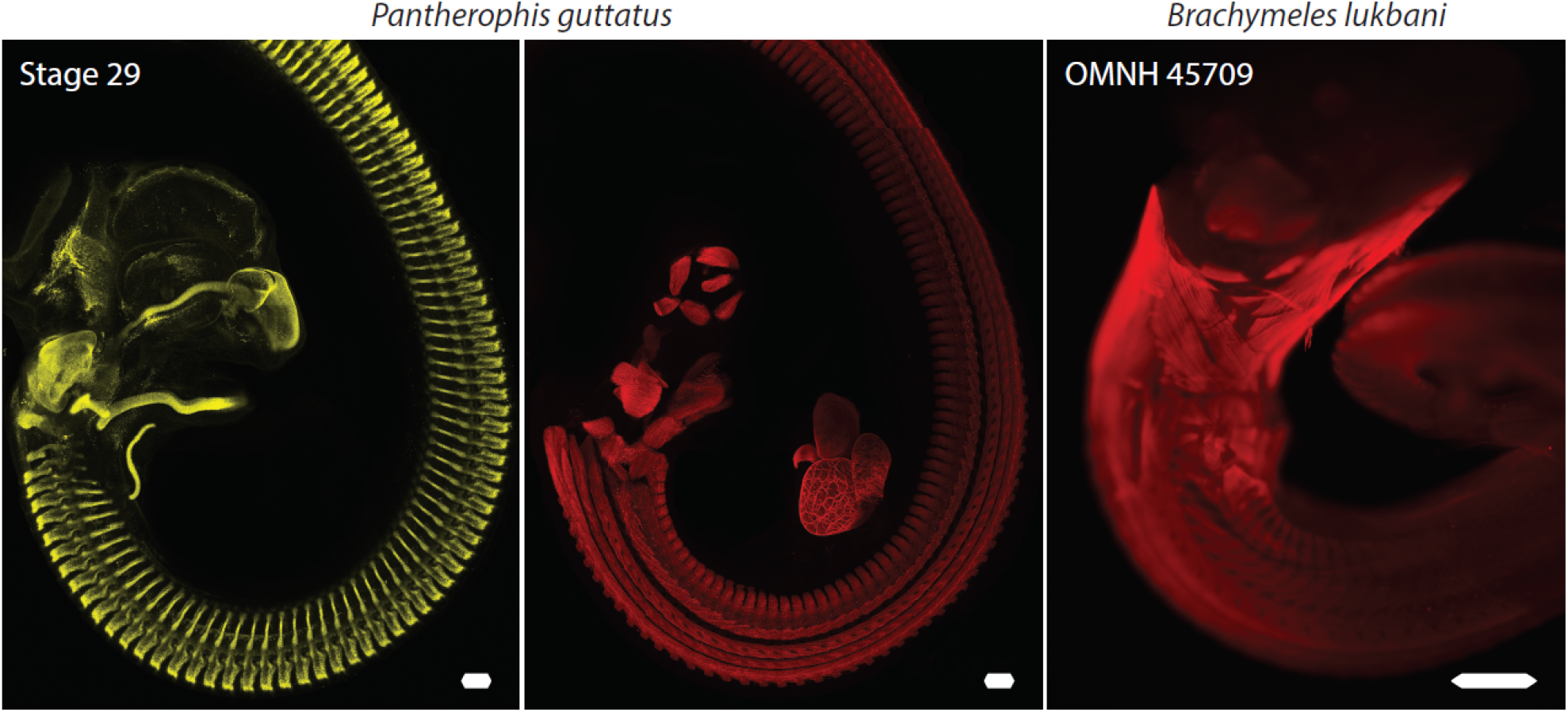
Comparison of the true limbless embryo of the corn snake, *Pantherophis guttatus*, with that of the limbed *Brachymeles lukbani*. Whereas *B. lukbani* develops both skeleton and muscles associated and dependent on the development of a limb bud, snakes show no trace of neither forelimb skeleton, girdles or musculature. Both the axial and the limb musculature of the skink, although reduced distally, develop in association with the pectoral girdle in the limb region, while the axial musculature of the snake remains undifferentiated along the anteroposterior axis

### 3.4 Comparison to embryos of the snake *Pantherophis guttatus*

In Stage 29 corn snake embryos, most of the skeleton development seems to correspond to the degree of development observed in *Brachymeles lukbani* embryo OMNH 45709, however, the postcranial skeleton shows no trace of pectoral girdle elements or limb skeleton (Figure 3). Axial musculature, in contrast to limb muscles, derives from muscles that develop first within the boundaries of the somite and then extend towards their specific attachments (Burke and Nowicki, 2003). None of the well-developed girdle axial muscles (see below) present in *B. lukbani* embryos can be observed in *Pantherophis.* Additionally, as expected in snakes, no trace of limb musculature, intrinsic or extrinsic, is observed in *Pantherophis* either.

### 3.5 Adult morphology of *Brachymeles lukbani*

Adult *B. lukbani* preserve a fairly well-developed, albeit thin and poorly ossified, pectoral girdle, a small and curved humerus, and extremely reduced radius and ulna (Figure 4 A, B). The pectoral girdle consists of a well-developed but undivided scapulocoracoid. The coracoid portion presents a well-defined metacoracoid only, with a primary and secondary coracoid ray delineating a primary coracoid fenestra (Russell and Bauer, 2008). The suprascapula is broad in its dorsal border, and probably calcified, as evidenced by the granular texture observed in the CTscan. The lateral two-thirds of the clavicles are heavily curved, and the medial ends are fenestrated. The interclavicle is arrow shaped, with an anterior process longer than the posterior process and lateral processes arching anteriorly. The sternum is calcified and bears a sternal fontanelle on its posterior end. The humerus is short and curved, with a pronounced humeral crest (Figure 4 A, B, E). Both the radius and ulna are extremely reduced, each a tiny splint of bone bone a few tens of microns in diameter (Figure 4 C, D). We confirmed that adult *B. lukbani* still possess well developed limb musculature associated with the pectoral girdle (Figure 4 F, G), such as broad latissimus dorsi and pectoralis muscles. The Deltoid musculature, on the other hand, while easily divisible into its scapular and clavicular portions in the embryos, is not so readily separated in the adult. The supracoracoid muscle, as in the embryos, has two, well defined portions originating from the coracoids and clavicles. A coracobrachialis muscle was identified, however other muscles deriving from the biceps, triceps, or more distal subdivisions were not observed.

**FIGURE 4.**
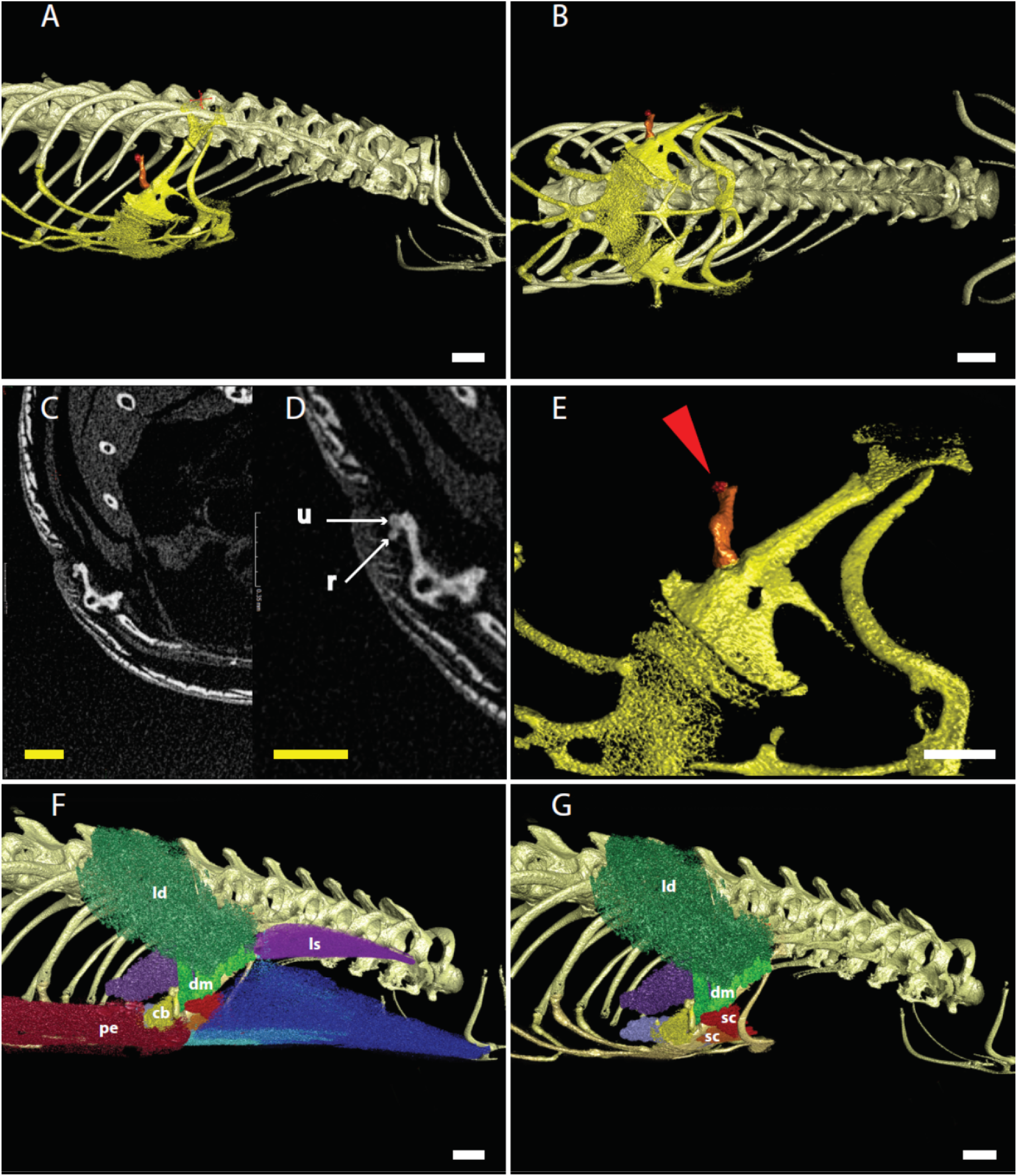
Adult morphology of *Brachymeles lukbani* obtained from μCT-scan imaging. A: Lateral view of the neck and thoracic region. B: Ventral view of the neck and thoracic region. The girdle elements are colored in yellow, humerus in orange, extremely reduced ulna and radius in red. C, D: CTscan raw data slices showing the humerus and the extremely reduced ulna and radius. E: Detail of the limb elements; humerus in orange, ulna and radius in red, pointed by the red arrow. F, G: Some of the muscles of the pectoral region, with limb muscles colored in green, red and yellow colors, and axial muscles in blue and purple colors. cb: coracobrachialis, dm: deltoid musculature, ld: latissimus dorsi, ls: levator scapulae, pe: pectoralis, r: radius, sc: supracoracoideus, u: ulna. White scale bars: 500 μm. Scale bar in C and D: 350 μm

## 4 DISCUSSION

It has been an enduring question is to what extent convergent limb reduction is achieved by employing the same developmental mechanisms. Limbs can be lost without losing the corresponding girdle, which probably indicates limbs are easier to lose or truncate during embryonic development than girdles. Across most clades of lizards, absence or reduction of forelimbs and/or pectoral girdles is not tied to reduction or loss of hindlimbs and/or pelvic girdles (Stephenson, 1962; Rodrigues, 1991; Nussbaum and Raxworthy, 1995; Pellegrino *et al*., 2001; Andreone and Greer, 2002; Sakata and Hikida, 2003a; Sakata and Hikida, 2003b; Rodrigues *et al*., 2008; Miralles *et al*., 2012). The prominent exception is of course the absence of forelimb elements in snakes. Otherwise, all “limbless” lizards for which embryonic development has been investigated develop limbs in early developmental stages, which later shrink, reabsorb or degenerate (Raynaud, 1985). As such, snakes are the only group of squamates studied in which adult true limblessness (fore- and hindlimbs absent in most snakes, only forelimbs absent in Aniliidae, Boidae, Leptotyphlopidae, Pythonidae and Typhlopidae,) reflects total absence of limb development in embryonic stages (Zehr, 1962; Raynaud, 1985; Jackson, 2002). Although the exact developmental mechanisms of each studied case are not completely understood, all seem to involve absent, reduced, or degenerated AER development or activity. In *Brachymeles lukbani*, the earliest forelimb observed displays a bent elbow and, although the autopod portion does not look properly developed, appears to be at a stage of development similar to when digit rays begin to develop in other lizards (Sanger *et al.*, 2008; Wise *et al.*, 2009; Rapp Py☐Daniel *et al.*, 2017; Griffing *et al.*, 2019). It was not possible to determine whether or not the AER of the limbs is normal in *B. lukbani* given available material; however, the earliest embryo seems to be around the temporal frame when the AER begins to become reduced in typical pentadactyl lizards, such as is observed in *Lacerta* (Raynaud, 2003) or *Paroedura* (*Noro et al*., 2009).

In amniote embryos, induction of early limb buds is, at least in part, dependent on signaling between the somites and the lateral plate mesoderm (LPM; Duester, 2008; Zhao *et al*., 2009; Zeller et al., 2009; Duboc and Logan, 2011), while the maintenance of limb development depends on the activity of the AER (Mahmood *et al.*, 1995). The forelimb skeleton develops from the mesoderm within the limb bud, derived from the LPM, while the pectoral girdle derives from the LPM and an additional component of somitic origin (McGonnell, 2001). The musculature of the forelimb and that connecting the limb skeleton to the pectoral girdle, and the girdle to the vertebral column, derives from the somites. During development, some cells migrate out of the somite and invade the limb buds where they differentiate into muscle cells and arrange forming two opposing muscle masses that give rise to the muscles of the arm (intrinsic limb muscles) and some major muscles originating on the pectoral girdle and extending to attachment points within the arm or on the axial column (extrinsic limb muscles) (i.e. pectoralis, supracoracoideus, latissimus, deltoideus, scapulohumeral, subscapular muscles). Other muscles form within the somite boundaries and extend into the girdles from their origin sites at the vertebrae or ribs (i.e. levator scapulae, trapezius, serratus, episternocleidomastoid muscles). The former group of muscles correspond to proper limb muscles, irrespective of their origin or attachments, as developmentally they derive from the limb muscle masses, while the latter group corresponds to axial musculature, as they originate developmentally from the somitic primaxial musculature (Romer, 1944; Russell and Bauer, 2008; Valasek *et al.*, 2011).

In the *limbless* chicken mutant, small limb buds start development but grow very little and soon after shrink and disappear (Prahlad *et al.*, 1979). In these mutant embryos, both the pectoral and pelvic girdles develop normally, however the limb skeletal elements and the limb musculature do not (Prahlad *et al.*, 1979). Furthermore, there is no sign of the humerus or more distal elements, nor of extrinsic limb muscles like the pectoralis, although axial girdle musculature appears to be normal (Prahlad *et al.*, 1979; Lanser and Fallon, 1984). This demonstrates that the maintenance of a developing limb bud is necessary for the formation and development of the limb skeleton and limb intrinsic and extrinsic musculature, at least up to a certain point, but is not needed for the development of the girdles nor the axial girdle musculature. In *Brachymeles lukani*, although development of the limb is truncated, its early presence seems to be sufficient to enable the development of the limb musculature. Subsequent limb reduction to the point of near-disappearance does not seem to affect the later development of extrinsic limb musculature associated with the normally developed pectoral girdle. This observation suggests that in other limbless clades, the presence of limb musculature in adults implies the presence of transient limbs during embryonic development.

As mentioned before, most lizard clades have evolved extremely limb-reduced or limbless forms. In fact; gekkotans, gerrhosaurids, cordylids, gymnophthalmids and anguimorphs display both limbed and limb-reduced or limbless species, and only in iguanians, lacertids, teiids and xantusids is limb loss not observed. Dibamids, amphisbaenians, and snakes are composed entirely of limb-reduced or limbless species (Figure 5). However, information on the developmental patterns and adult muscle anatomy of limbs and girdles remains scarce. Amphisbaenians form a highly specialized fossorial clade of lizards, composed by five families characterized by limblessness (Kearney, 2002). Only members of the genera *Blanus* and *Bipes* retain a reduced femur, and only species of *Bipes* have forelimbs, which are well developed and include humerus, ulna, radius, carpals, and four or five digits (Kearney, 2002). In stark contrast to *Bipes*, all other amphisbaenians lack any trace of forelimb skeletal elements. However, with the exception of the family Rhineuridae, all have been reported recently to retain the ancestral number of forelimb girdle muscles, although these muscles show somewhat modified arrangements, and origin and attachment points, associated with their variably developed pectoral girdles (Westphal *et al.*, 2019). Interestingly, Rhineuridae also lacks any pectoral girdle skeletal element, but does possess highly modified strand-like muscles that are similar to those of other amphisbaenians and lizards in position and number (Westphal *et al*., 2019). The presence of axial pectoral and limb girdle musculature in amphisbaenians suggests they may retain a developing forelimb, at least during early embryonic stages. The retention of an early forelimb during development may explain the apparent re-evolution of forelimbs or digits without the necessity of invoking novel re-evolution of limb development mechanisms and processes in an ancestrally limbless clade, not only in the case of *Bipes biporus* (Kearney and Stuart, 2004; Brandley *et al*., 2008), but also in analogous cases within Gymnophthalmidae (Kohlsdorf and Wagner, 2006) and Scincidae (Wagner *et al.*, 2018).

**FIGURE 5.**
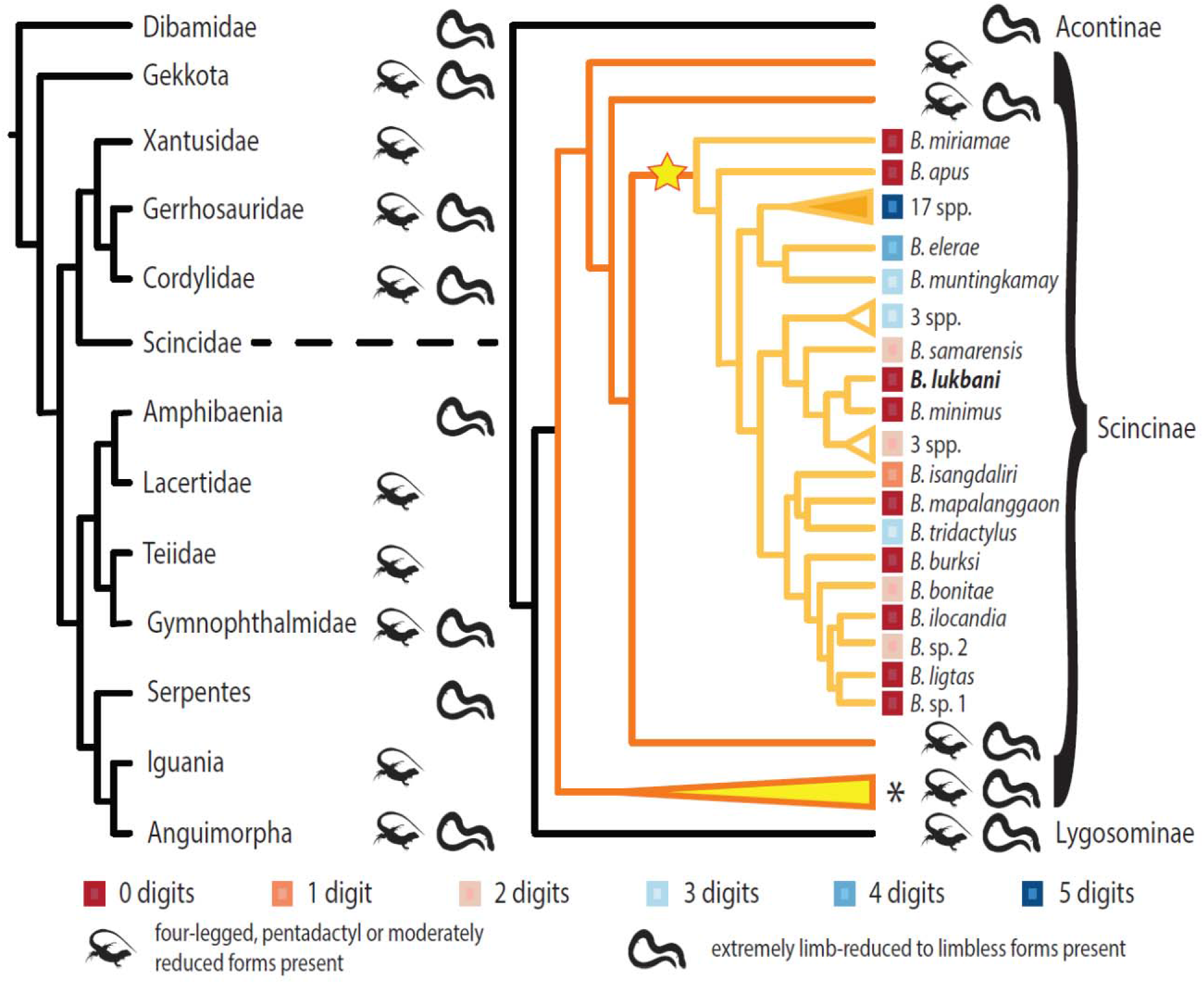
Phylogenetic relationships among skinks in the genus *Brachymeles* in the context of lizard phylogeny and limb reduction. Extreme limb-reduction has evolved independently within most clades of squamates, with the exception of Xantusidae, Lacertidae, Teiidae and Iguania, while Dibamidae, Amphisbaenia and Serpentes are composed exclusively by extremely limb-reduced or limbless species. Whithin Scincidae (Right), Acontinae is composed exclusively of extremely limb-reduced or limbless forms, and limbs have ben reduced or lost many times within Lygosominae and Scincinae. Nested within Scincinae (orange branches of the tree, the genus *Brachymeles* (light orange branches, node marked by a star) displays an interesting pattern of multiple independent events of extreme limb reduction, exemplified by the number of digits retained in the forelimb (colored squares). Orange colored triangle represents a *Brachymeles* lineage composed of 17 pentadactyl species. White colored trianlges represent lineages of *Brachymeles* with three species of similar degrees of limb reduction (2 or 3 digits retained), and the yellow colored tringles represents the rest of genera whithin Scincinae, including at least 20 genera with extremely limb reduced species (*Chalcides*, *Scelotes*, *Feylinia*, *Jarujinia*, *Pygomeles*, among others, for example). Squamate tree modified from Leal and Cohn, 2018. Skink phylogeny modified from Pyron *et al*., 2013; Andrade *et al.*, 2016; Wagner *et al.*, 2018

Skinks are without a doubt the best clade to study the evolution of limb reduction and loss among squamates. Extreme limb reduction and limblessness is observed in species in more than 30 different genera representing an even larger number of independent transitions from the pentadactyl ancestral state. As in many skink clades, instances of limb reduction and loss occur independently in lineages of the same genus, as is observed in *Brachymeles* (Figure 5). *Brachymeles lukbani* has an at least externally limbless sister species, *B. minimus*, and both are nested in a clade of seven species with reduced limb but digited lineages (Figure 5; Wagner *et al*., 2018; Bergmann *et al*., 2020). Within this clade, the two-digited forms are more closely related to *B. lukbani* and *B. minimus*, sister to the three-digited species (Figure 5). This pattern suggests a progressive loss of digits and more proximal limb structures in the *lukbani* + *minimus* clade. Based on a dated phylogeny (Wagner *et al.*, 2018; Bergmann *et al.*, 2020) limb loss in these animals is relatively recent, probably less than 12 million years ago. This phylogenetic history is consistent with a developmental pattern that still includes the embryonic appearance of stylo and zeugopodium and shoulder girdle and associated muscles, and retention of a reduced limb covered under a scale, a condition called *cyptomelia* (Windle, 1898).

Within individual genera, the morphology of reduced limbs and the degree of reduction can be highly variable (Greer, 1970; Andreone and Greer, 2002; Sakata and Hikida, 2003a; Sakata and Hikida, 2003b; Carranza *et al*., 2008; Siler *et al*., 2011a; Davis *et al*., 2014; Miralles *et al*., 2015; Andrade *et al*., 2016; Wagner *et al*., 2018). The persistence of intermediate forms between the fully pentadactyl and fully limbless represents a mystery, that has been interpreted as evidence that these species represent different adaptive optima (Brandley *et al*., 2008; Bergmann and Morinaga, 2019; Skinner *et al*., 2008; Bergmann and Morinaga, 2019), although optimal for what is not known. The extent to which extremely reduced rudimentary limbs and their associated muscles participate actively in locomotion is largely unknown (Bergmann *et al.*, 2020). There are indications that different morphologies do not affect locomotor performance (Morinaga and Bergmann, 2020), further conflicting with the notion that intermediate forms are adaptive (but see Bergmann *et al.*, 2020). The high variability in digit number and degree of reduction seen at the interspecific level mirrored even within some individual species (Siler *et al*., 2011b; Davis *et al*., 2014; Andrade *et al*., 2016). This parallel of morphological variability between species in a genus and among individuals within a species suggests that drift, constrained by population sizes and isolation, rather than that active adaptation plays an important role in the maintenance of intermediate limb-reduced morphologies. Meanwhile, the apparent progression from moderate to extreme limb reduction observed in different limb reduced lineages might hint at cumulative developmental effects behind the initial evolution and persistence of reduced limb morphologies.

It is worth noting that the situation in *B. lukbani* is unusual as this species is externally limbless but retains a hidden limb rudiment, a condition that has been called *cryptomelia* in the medical literature (Windle, 1898). It is not clear how common this form of limbless phenotype is, as it requires special techniques to detect. Cryptic limbs like these may have gone undetected in many other apparently limbless lizard species. The results of this study provide a reasonable scenario linking the transient existence of a developing limb with the presence of limb derived girdle muscles in adults. Nevertheless, studies on other limb reduced taxa are needed to establish whether transient embryonic limbs or cryptic adult limbs are regularly present in species where these muscles are well developed but show no traces of limb skeleton. Further studies comparing the adult musculoskeletal anatomy, embryonic development, and phylogenetic evolutionary patterns of limb reduction in other skinks in the genus *Brachymeles*, as well as in the many other limb-reduced lineages, are required to understand the recurrent evolution of limb reduced forms in squamates and whether these similar phenotypes are the result of similar mechanisms evolving in parallel.

## ACKNOWLEDGEMENTS

We are particularly grateful to Mr. Jason Fernandez and our Filipino colleagues who were instrumental in carrying out successful field expeditions throughout this work. We would also like to thank Brandon Mercado for assistance with μCT-scanning. We thank the Sam Noble Oklahoma Museum of Natural History for granting access to its specimens. This research was supported by the Yale Institute of Biospheric Studies at Yale University, the Peabody Museum of Natural History, and the following NSF grants: 1353683, 1353743, 1353691, and 1353703.

## AUTHOR CONTRIBUTIONS

Oliver Griffith, Cameron D. Siler and Gunter P. Wagner collected the *Brachymeles lukbani* embryos and adults on the field. Matteo Fabbri and Laurel Yohe mounted and CT-scanned the adult specimens. Daniel Blackburn provided corn snake eggs from his colony at Trinity College. Bhart-Anjan S. Bhullar provided logistical and financial support for the immunostaining and microscopic imaging of the embryos. Daniel Smith-Paredes conceived the study, photographed *B. lukbani embryos*, collected corn snake embryos, performed the immunostaining experiments, imaged the immunostained embryos, created the figures and wrote manuscript with the assistance of Cameron D. Siler and Gunter P. Wagner.

## DATA AVAILABILITY STATEMENT

The data that support the findings of this study are available from the corresponding author upon reasonable request

## REFERENCES

Andrade, J. B., Lewis, R. P. & Senter, P. 2016. Appendicular skeletons of five Asian skink species of the genera *Brachymeles* and *Ophiomorus*, including species with vestigial appendicular structures. Amphibia-Reptilia, 37(4), pp 337–344.

Andreone, F. & Greer, A. E. 2002. Malagasy scincid lizards: descriptions of nine new species, with notes on the morphology, reproduction and taxonomy of some previously described species (Reptilia, Squamata: Scincidae). Journal of Zoology, 258(2), pp 139–181.

Bergmann, P. J. & Morinaga, G. 2019. The convergent evolution of snake-like forms by divergent evolutionary pathways in squamate reptiles. Evolution, 73(3), pp 481–496.

Bergmann, P. J., Morinaga, G., Freitas, E. S., Irschick, D. J., Wagner, G. P. & Siler, C. D. 2020. Locomotion and palaeoclimate explain the re-evolution of quadrupedal body form in *Brachymeles* lizards. Proceedings of the Royal Society B, 287(1938), pp 20201994.

Brandley, M. C., Huelsenbeck, J. P. & Wiens, J. J. 2008. Rates and patterns in the evolution of snake-like body form in Squamate reptiles: Evidence for repeated re-evolution of lost digits and long-term persistence of intermediate body forms. Evolution, 62(8), pp 2042–2064.

Burke, A. C. & Nowicki, J. 2003. A new view of patterning domains in the vertebrate mesoderm. Developmental Cell, 4(2), pp 159–165.

Carranza, S., Arnold, E., Geniez, P., Roca, J. & Mateo, J. 2008. Radiation, multiple dispersal and parallelism in the skinks, *Chalcides* and *Sphenops* (Squamata: Scincidae), with comments on *Scincus* and *Scincopus* and the age of the Sahara Desert. Molecular Phylogenetics and Evolution, 46(3), pp 1071–1094.

Christ, B. & Brand-Saberi, B. 2004. Limb muscle development. International Journal of Developmental Biology, 46(7), pp 905–914.

Cohn, M. J. & Tickle, C. 1999. Developmental basis of limblessness and axial patterning in snakes. Nature, 399(6735), pp 474.

Davis, D. R., Feller, K. D., Brown, R. M. & Siler, C. D. 2014. Evaluating the diversity of Philippine slender skinks of the *Brachymeles bonitae* Complex (Reptilia: Squamata: Scincidae): redescription of B. tridactylus and descriptions of two new species. Journal of Herpetology, 48(4), pp 480–494.

Duboc, V. & Logan, M. P. 2011. Regulation of limb bud initiation and limb-type morphology. Developmental Dynamics, 240(5), pp 1017–1027.

Duester, G. 2008. Retinoic acid synthesis and signaling during early organogenesis. Cell, 134(6), pp 921–931.

Gans, C. 1975. Tetrapod limblessness: evolution and functional corollaries. American Zoologist, 15(2), pp 455–467.

Greer, A. E. 1970. The Systematics and Evolution of the Subsaharan Africa, Seychelles, and Mauritius Scincine Scincid Lizards. Bull. Mus. Comp. Zool., 140 (1-23).

Greer, A. E. 1991. Limb reduction in squamates: identification of the lineages and discussion of the trends. Journal of Herpetology, 166–173.

Griffing, A. H., Sanger, T. J., Daza, J. D., Nielsen, S. V., Pinto, B. J., Stanley, E. L. & Gamble, T. 2019. Embryonic development of a parthenogenetic vertebrate, the mourning gecko (*Lepidodactylus lugubris*). Developmental Dynamics, 248(11), pp 1070–1090.

Infante, C. R., Rasys, A. M. & Menke, D. B. 2018. Appendages and gene regulatory networks: Lessons from the limbless. genesis, 56(1), pp e23078.

Jackson, K. 2002. Post-ovipositional development of the monocled cobra, *Naja kaouthia* (Serpentes: Elapidae). Zoology, 105(3), pp 203–214.

Kearney, M. 2002. Appendicular skeleton in amphisbaenians (Reptilia: Squamata). Copeia, 2002(3), pp 719–738.

Kearney, M. & Stuart, B. L. 2004. Repeated evolution of limblessness and digging heads in worm lizards revealed by DNA from old bones. Proceedings of the Royal Society of London. Series B: Biological Sciences, 271(1549), pp 1677–1683.

Kohlsdorf, T. & Wagner, G. P. 2006. Evidence for the reversibility of digit loss: a phylogenetic study of limb evolution in *Bachia* (Gymnophthalmidae: Squamata). Evolution, 60(9), pp 1896–1912.

Lande, R. 1978. Evolutionary mechanisms of limb loss in tetrapods. Evolution, 73–92.

Lanser, M. E. & Fallon, J. F. 1984. Development of the lateral motor column in the *limbless* mutant chick embryo. Journal of Neuroscience, 4(8), pp 2043–2050.

Leal, F. & Cohn, M. J. 2018. Developmental, genetic, and genomic insights into the evolutionary loss of limbs in snakes. Genesis, 56(1), pp e23077.

List, J. C. 1966. Comparative osteology of the snake families Typhlopidae and Leptotyphlopidae. 36. Illinois biological monographs; v. 36.

Mahmood, R., Bresnick, J., Hornbruch, A., Mahony, C., Morton, N., Colquhoun, K., Martin, P., Lumsden, A., Dickson, C. & Mason, I. 1995. A role for FGF-8 in the initiation and maintenance of vertebrate limb bud outgrowth. Current biology, 5(7), pp 797–806.

McGonnell, I. M. 2001. The evolution of the pectoral girdle. Journal of Anatomy, 199(1-2), pp 189–194.

Miralles, A., Anjeriniaina, M., Hipsley, C. A., Müller, J., Glaw, F. & Vences, M. 2012. Variations on a bauplan: description of a new Malagasy “mermaid skink” with flipper-like forelimbs only (Scincidae, *Sirenoscincus* Sakata & Hikida, 2003). Zoosystema, 34(4), pp 701–720.

Miralles, A., Hipsley, C. A., Erens, J., Gehara, M., Rakotoarison, A., Glaw, F., Müller, J. & Vences, M. 2015. Distinct patterns of desynchronized limb regression in Malagasy scincine lizards (Squamata, Scincidae). PLoS One, 10(6), pp e0126074.

Morinaga, G. & Bergmann, P. J. 2020. Evolution of fossorial locomotion in the transition from tetrapod to snake-like in lizards. Proceedings of the Royal Society B, 287(1923), pp 20200192.

Noro, M., Uejima, A., Abe, G., Manabe, M. & Tamura, K. 2009. Normal developmental stages of the Madagascar ground gecko *Paroedura pictus* with special reference to limb morphogenesis. Developmental dynamics: an official publication of the American Association of Anatomists, 238(1), pp 100–109.

Nussbaum, R. A. & Raxworthy, C. J. 1995. Review of the scincine genus *Pseudoacontias* Barboza du Bocage (Reptilia: Squamata: Scincidae) of Madagascar. Herpetologica, 91–99.

Pellegrino, K. C., Rodrigues, M. T., Yonenaga-Yassuda, Y. & Sites Jr, J. W. 2001. A molecular perspective on the evolution of microteiid lizards (Squamata, Gymnophthalmidae), and a new classification for the family. Biological Journal of the Linnean Society, 74(3), pp 315–338.

Prahlad, K., Skala, G., Jones, D. G. & Briles, W. 1979. Limbless: a new genetic mutant in the chick. Journal of Experimental Zoology, 209(3), pp 427–434.

Pyron, R. A., Burbrink, F. T. & Wiens, J. J. 2013. A phylogeny and revised classification of Squamata, including 4161 species of lizards and snakes. BMC evolutionary biology, 13(1), pp 1–54.

Rahmanl, T. M.-Z. 1974. Morphogenesis of the rudimentary hind-limb of the Glass Snake (*Ophisaurus apodus* Pallas). Development, 32(2), pp 431–443.

Rapp Py-Daniel, t., Kennedy Soares De-Lima, A., Campos Lima, F., Pic-Taylor, A., Rodrigues Pires Junior, O. & Sebben, A. 2017. A staging table of post-ovipositional development for the South American collared lizard *Tropidurus torquatus* (Squamata: Tropiduridae). The Anatomical Record, 300(2), pp 277–290.

Raynaud, A. 1985. Development of limbs and embryonic limb reduction. Biology of the Reptilia, 15(59-148).

Raynaud, A. 2003. Developmental mechanism involved in the embryonic reduction of limbs in reptiles. International Journal of Developmental Biology, 34(1), pp 233–243.

Raynaud, A., Gasc, J. & Renous-Lecuru, S. 1975. Les rudiments de membres et leur développement embryonnaire chez *Scelotes inornatus* (A. Smith)(Scincidae, Sauria). Bull Mus Nat Hist Nat (Paris), 208(537-551).

Raynaud, A. & Van den Elzen, P. 1976. La rudimentation des membres chez les embryons de *Scelotes gronovii* (Daudin), reptile scincidé Sud-Africain. Arch. Anat. Microsc. Morphol. Exp, 65(17-36).

Rodrigues, M. T. 1991. Herpetofauna das dunas interiores do Rio Sao Francisco, Bahia, Brasil. 1: Introducao a área e descricao de um novo género de Microteiideos (*Calyptommatus*) com notas sobre sua ecologia, distribuicao e especiacao (Sauria, teiidae). Papeis Avulsos de Zoologia, Sao Paulo, 27(329-342).

Rodrigues, M. T., Camacho, A., Nunes, P. M. S., Recoder, R. S., Teixeira Jr, M., Valdujo, P. H., Ghellere, J. M. B., Mott, T. & Nogueira, C. 2008. A new species of the lizard genus *Bachia* (Squamata: Gymnophthalmidae) from the Cerrados of Central Brazil. Zootaxa, 1875(1), pp 39–50.

Romer, A. S. 1944. The development of tetrapod limb musculature—the shoulder region of *Lacerta*. Journal of Morphology, 74(1), pp 1–41.

Russell, A. & Bauer, A. M. 2008. The appendicular locomotor apparatus of *Sphenodon* and normal-limbed squamates. *In*: C. Gans, A. G. K. A. (ed.) Biology of the Reptilia. Ithaca, NY: Society for the Study of Amphibians and Reptiles.

Sakata, S. & Hikida, T. 2003a. A Fossorial Lizard with Forelimbs Only. Current herpetology, 22(1), pp 9–15.

Sakata, S. & Hikida, T. 2003b. A new fossorial scincine lizard of the genus *Pseudoacontias* (Reptilia: Squamata: Scincidae) from Nosy Be, Madagascar. Amphibia-Reptilia, 24(1), pp 57–64.

Sanger, T. J., Losos, J. B. & Gibson-Brown, J. J. 2008. A developmental staging series for the lizard genus *Anolis*: a new system for the integration of evolution, development, and ecology. Journal of Morphology, 269(2), pp 129–137.

Schindelin, J., Arganda-Carreras, I., Frise, E., Kaynig, V., Longair, M., Pietzsch, T., Preibisch, S., Rueden, C., Saalfeld, S. & Schmid, B. 2012. Fiji: an open-source platform for biological-image analysis. Nature methods, 9(7), pp 676.

Siler, C. D., Balete, D. S., Diesmos, A. C. & Brown, R. M. 2010. A new legless loam-swimming lizard (Reptilia: Squamata: Scincidae: Genus *Brachymeles*) from the Bicol Peninsula, Luzon Island, Philippines. Copeia, 2010(1), pp 114–122.

Siler, C. D. & Brown, R. M. 2011. Evidence for repeated acquisition and loss of complex body-form characters in an insular clade of Southeast Asian semi-fossorial skinks. Evolution: International Journal of Organic Evolution, 65(9), pp 2641–2663.

Siler, C. D., Diesmos, A. C., Alcala, A. C. & Brown, R. M. 2011a. Phylogeny of Philippine slender skinks (Scincidae: *Brachymeles*) reveals underestimated species diversity, complex biogeographical relationships, and cryptic patterns of lineage diversification. Molecular Phylogenetics and Evolution, 59(1), pp 53–65.

Siler, C. D., Fuiten, A. M., Jones, R. M., Alcala, A. C. & Brown, R. M. 2011b. Phylogeny-based species delimitation in Philippine slender skinks (Reptilia: Squamata: Scincidae) II: taxonomic revision of *Brachymeles samarensis* and description of five new species. Herpetological Monographs, 25(1), pp 76–112.

Simmons, J. 2015. Herpetological Collecting and Collections Management. Society for the Study of Amphibians and Reptiles Herpetology, Circular 42(1-210).

Skinner, A. & Lee, M. S. 2009. Body-form evolution in the scincid lizard clade *Lerista* and the mode of macroevolutionary transitions. Evolutionary Biology, 36(3), pp 292–300.

Skinner, A., Lee, M. S. & Hutchinson, M. N. 2008. Rapid and repeated limb loss in a clade of scincid lizards. BMC Evolutionary Biology, 8(1), pp 1–9.

Stephenson, N. 1962. The comparative morphology of the head skeleton, girdles and hind limbs in the Pygopodidae. Zoological Journal of the Linnean Society, 44(300), pp 627–644.

Valasek, P., Theis, S., DeLaurier, A., Hinits, Y., Luke, G. N., Otto, A. M., Minchin, J., He, L., Christ, B. & Brooks, G. 2011. Cellular and molecular investigations into the development of the pectoral girdle. Developmental biology, 357(1), pp 108–116.

Wagner, G. P., Griffith, O. W., Bergmann, P. J., Bello-Hellegouarch, G., Kohlsdorf, T., Bhullar, A. & Siler, C. D. 2018. Are there general laws for digit evolution in squamates? The loss and re-evolution of digits in a clade of fossorial lizards (*Brachymeles*, Scincinae). Journal of Morphology, 279(1104-19).

Westphal, N., Mahlow, K., Head, J. J. & Müller, J. 2019. Pectoral myology of limb-reduced worm lizards (Squamata, Amphisbaenia) suggests decoupling of the musculoskeletal system during the evolution of body elongation. BMC evolutionary biology, 19(1), pp 1–23.

Wiens, J. J. & Slingluff, J. L. 2001. How lizards turn into snakes: a phylogenetic analysis of body-form evolution in anguid lizards. Evolution, 55(11), pp 2303–2318.

Windle, B. C. 1898. Eighth Report on Recent Teratological Literature. Journal of Anatomy and Physiology, 32(Pt 4), pp 780.

Wise, P. A., Vickaryous, M. K. & Russell, A. P. 2009. An embryonic staging table for in ovo development of *Eublepharis macularius*, the leopard gecko. The Anatomical Record: Advances in Integrative Anatomy and Evolutionary Biology: Advances in Integrative Anatomy and Evolutionary Biology, 292(8), pp 1198–1212.

Yang, B., Treweek, J. B., Kulkarni, R. P., Deverman, B. E., Chen, C.-K., Lubeck, E., Shah, S., Cai, L. & Gradinaru, V. 2014. Single-cell phenotyping within transparent intact tissue through whole-body clearing. Cell, 158(4), pp 945–958.

Young, R. L., Caputo, V., Giovannotti, M., Kohlsdorf, T., Vargas, A. O., May, G. E. & Wagner, G. P. 2009. Evolution of digit identity in the three-toed Italian skink *Chalcides chalcides*: A new case of digit identity frame shift. Evolution & development, 11(6), pp 647–658.

Zehr, D. R. 1962. Stages in the normal development of the common garter snake, *Thamnophis sirtalis sirtalis*. Copeia, 322–329.

Zeller, R., López-Ríos, J. & Zuniga, A. 2009. Vertebrate limb bud development: moving towards integrative analysis of organogenesis. Nature Reviews Genetics, 10(12), pp 845.

Zhao, X., Sirbu, I. O., Mic, F. A., Molotkova, N., Molotkov, A., Kumar, S. & Duester, G. 2009. Retinoic acid promotes limb induction through effects on body axis extension but is unnecessary for limb patterning. Current Biology, 19(12), pp 1050–1057.

